# Preliminary stability studies of a ß-SARS-CoV-2 virus-like particle vaccine

**DOI:** 10.64898/2026.03.11.711036

**Authors:** Melissa A. Edeling, Linda Earnest, Julio Carrera Montoya, Ashley Huey Yiing Yap, Dhiraj Hans, Joseph Torresi

## Abstract

We aimed to study the stability of a ß-SARS-CoV-2 virus-like particle (VLP) vaccine in a series of preliminary experiments using select stabilising excipients. ß-SARS-CoV-2 VLPs were produced and purified using established methodologies. The thermostability of VLPs was tested at 4°C and -30°C in the presence or absence of stabilizers polysorbate 80, sorbitol or L-histidine in the presence of a physiological NaCl concentration of 137mM. The integrity of VLPs was assessed using ELISA, Western immunoblot and dynamic light scatter (DLS). ß-SARS-CoV-2 VLPs were stable at 4°C for 14 days and the addition of stabilizing excipients improved stability compared to VLPs in PBS alone. Storage of VLPs at -80°C maintained particle integrity by DLS analysis for up to 2 years. Excipients helped to maintain the immunogenicity of the VLPs by ELISA and Western immunoblot and DLS analysis revealed that VLPs retained their particulate structure.

**Importance:** SARS-CoV-2 continues to circulate globally and cause significant illness. The problem of waning immunity to mRNA/LNPs has necessitated frequent boosters to keep pace of emerging variants. The development of alternative vaccines therefore remans a priority. Protein based vaccines, like VLPs, offer a safe alternative able to produce longer lasting immune responses. In this preliminary stability analysis, the ß-SARS-CoV-2 VLPs were found to be stable at 4°C and the addition of excipients improved VLP stability. Storage of VLPs at -30°C and -80°C also showed that the VLPs are stable for very long periods. Our findings will be of importance for the ongoing development of a SARS-CoV-2 VLP based vaccine.

## 1. Introduction

The COVID-19 pandemic led to the development of several vaccines based on different platforms including mRNA/LNPs, recombinant adenoviral vectored vaccines, inactivated viral vaccines, subunit and nanoparticle vaccines. Whilst mRNA/LNPs became the most widely used globally and had significant impacts in helping to control the pandemic, they suffered from short lived immune responses, thereby necessitating frequent boosters and re-development of variant specific mRNA/LNPs.

Virus like particle (VLP) vaccines for hepatitis B, human papilloma virus and hepatitis E virus have been shown to be highly effective (1-3). The repetitive, ordered, particulate structure of VLPs makes these an attractive vaccine candidate for enveloped RNA viruses, although to date there are no commercial VLP vaccines for these viruses (4-8). A further advantage of VLP based vaccines is that they can stimulate the production of protective CD4+ and CD8+ T cell responses in addition to protective antibody responses.

We have developed a ß-SARS-CoV-2 VLP vaccine from a single self-cleaving polyprotein and that includes the key structural proteins spike (S), envelope (E) and membrane (M), which are required to assemble authentic VLPs (9) (10-12). Using industry styled methods, it was possible to scale up production and purification of the ß-SARS-CoV-2 VLPs providing reproducible batches of vaccine (11). Formulating the purified ß-SARS-CoV-2 VLPs with MF59 produced strong and protective immune responses in mice against homologous and heterologous viral challenge (13).

For ß-SARS-CoV-2 VLPs to be used as a vaccine it is important to maintain the integrity of the VLPs during long-term storage, as a change in their conformational structure could affect vaccine potency and make the VLPs less effective in providing protection against SARS-CoV-2 variants. To maintain the stability and structural integrity of the VLPs it is necessary to include protective molecules in the final formulation buffer including varying combinations and concentrations of excipients like trehalose, sucrose, polysorbate 80, sorbitol and amino acids like L-histidine, arginine, glycine and proline (14-16). These stabilisers protect VLPs from degradation during freeze-thawing and prevent VLP aggregation that could result in loss of vaccine potency and batch to batch variation in vaccine quality that could impact on vaccine licensure.

Characterisation of the VLPs through the course of stability studies also requires a panel of well validated assays, including ELISA, Western blot and a measure of particle size and integrity (17) (14, 18). Dynamic light scatter (DLS) is frequently used to determine particle size and integrity because of its ease of use and access (15, 18, 19). However, more accurate analysis of size, size distribution, degradation and aggregation of VLPs that can meet industry standards can be achieved by combining DLS with methods like asymmetric flow field fractionation (AF4) and multiangle light scatter (15, 20).

In this study we evaluated the stability of ß-SARS-CoV-2 VLPs under different conditions with three excipients, polysorbate 80, sorbitol and L-histidine. The VLPs were analysed by ELISA, Western immunoblot and dynamic light scatter (DLS) and were found to be stable at 4°C and to maintain their particulate integrity.

## 2. Methods

### 2.1 Production and purification of ß-SEM-SARS-CoV-2 VLPs

The method for producing, purifying and titrating the recombinant adenoviruses expressing SARS-CoV-2 S, E and M genes has been described previously (9, 10, 21). The recombinant adenovirus was used to infect Vero cells in cell factories to produce ß-SEM-SARS-CoV-2 VLPs as described previously (9, 10, 21). In brief, VLPs were produced using 5 stack cell factories (Corning Life Sciences, USA). Vero cells (4.9x10^6^) were seeded into each 175 cm^2^ layer (2.45x10^7^ cells per 5-stack cell factory) in 20ml per layer of OPTIPRO SFM (Gibco, Thermofisher Scientific, USA) supplemented with 2% Glutamax (Gibco, Thermofisher Scientific, USA), 1% penicillin and streptomycin and incubated overnight at 37°C in 4% CO_2_. Vero cells at 80% confluency (18.6 x 10^6^ cells) were infected with recombinant adenovirus at an MOI of 1.0. ß-SEM-SARS-CoV-2 VLPs were purified from cell culture supernatants volume using a sequential process of tangential flow filtration (TFF), diafiltration, anion exchange chromatography, ultrafiltration and finally sterilized through a 0.45µm filter (11, 21).

### 2.2 Western immunoblot

Following separation by SDS-PAGE and transfer to polyvinylidene fluoride (PVDF) membrane, blots were probed with anti-S (40591-T62, Sino Biologicals), anti-M (MBS434281, MyBioSource) and anti-E (MBS8309656, MyBioSource) antibodies for SARS-CoV-2 VLPs as described previously (11, 21).

### 2.3 Enzyme Linked Immunosorbent Assay (ELISA)

Spike protein was assayed using ELISA by directly coating a Nunc MaxiSorp™ flat-bottom 96 well plate (Cat # NUN439454) overnight with VLPs at a concentration of 20 µg/mL and probing with anti-Spike detection antibody (polyclonal rabbit antibody Sinobiological 40591-T62) as described previously (9, 11, 21).

### 2.4 Dynamic Light Scattering (DLS)

Light scattering intensity fluctuations of purified SARS-CoV-2 VLP samples were measured on a Zetasizer Ultra (Malvern Panalytical) using ZS Xplorer software as described previously (11). Samples were incubated at 4°C and DLS measurements were collected over 15 cycles, in triplicate.

## 3. Results

### 3.1 Production, purification and characterization of ß-SARS-CoV-2 VLPs

We have previously described the production of ß-SARS-CoV-2 VLPs using a recombinant adenovirus vector carrying a gene construct containing the full-length spike (S) stabilized by the introduction of proline substitutions A892P and A942P (22), envelope (E) and membrane (M) genes (9, 11). Adenoviruses were used to transduce HEK 293A cells followed by passaging to produce a high titre viral stock that was subsequently used to infect Vero cell factories to produce ß-SARS-CoV-2 VLPs. The VLPs were purified using a process of clarification followed by tangential flow filtration, diafiltration, anion exchange chromatography and finally sterile ultrafiltration before aliquoting and storage at -80°C (11).

In a preliminary thermostability experiment, VLPs were stored without excipients at 4°C or - 30°C for 4 months and analysed by ELISA probing with anti-RBD antibody. Particles stored at -30°C retained stronger anti-RBD binding compared to particles stored at 4°C. However, VLPs stored at 4°C were still strongly reactive to anti-RBD antibody (Figure 1). VLPs stored at -80°C for 6 to 12 months retained anti-RBD and anti-S reactivity (results not shown).

**Figure 1.**
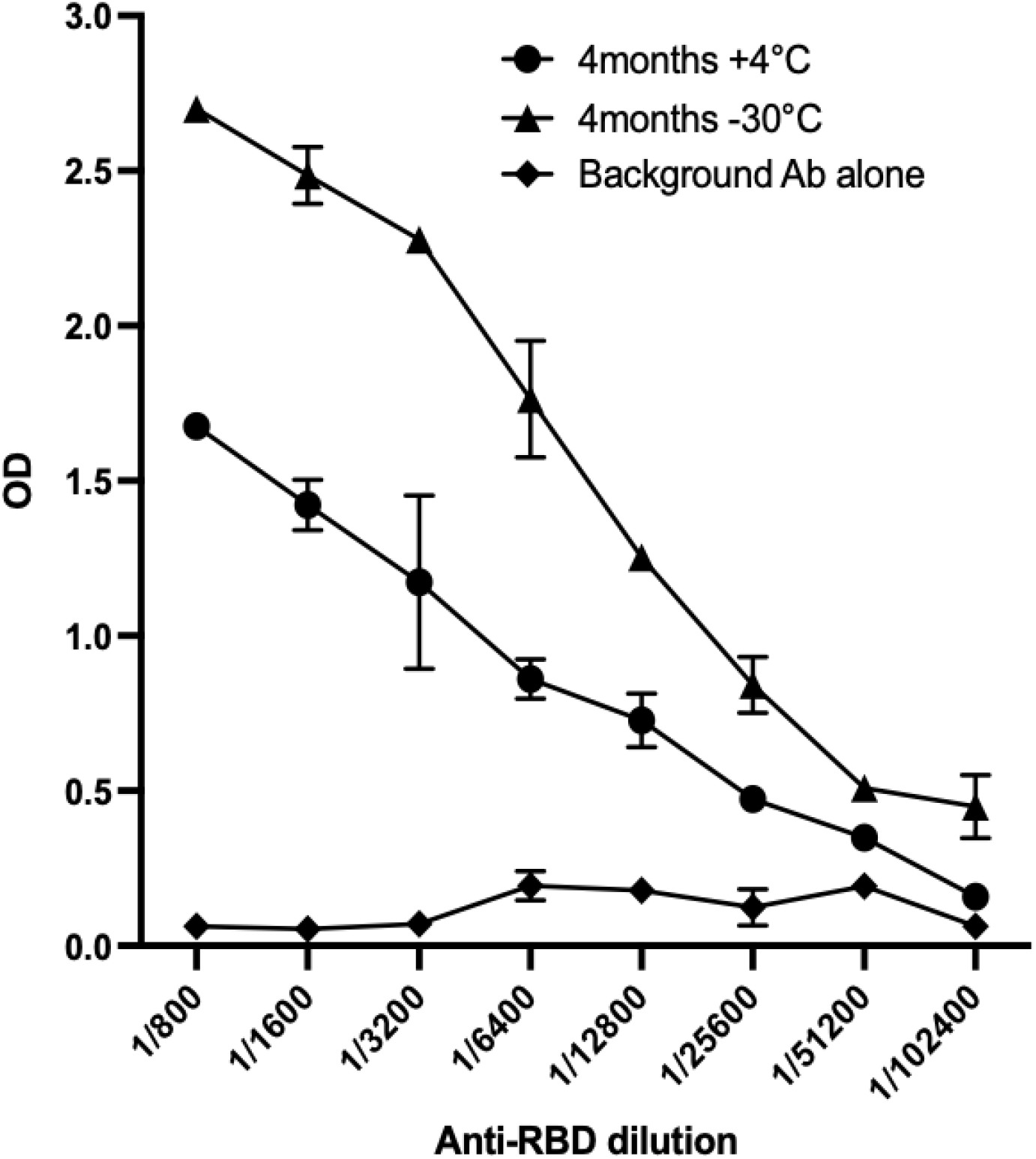
ELISA analysis of SARS-CoV-2 VLPs stored at 4°C or -30°C for 4 months. Plates were coated with SARS-CoV-2 VLPs followed by probing with anti-RBD antibody in 2-fold dilutions from 1:800 to 1:102,400 (mean ± SD). ELISAs were repeated three times and each dilution tested in triplicate.

### 3.2 ELISA of ß-SARS-CoV-2 VLPs with excipients

To determine whether the addition of different molecules, including salts, non-ionic surfactants, sugars or amino acids could act to stabilise the VLPs, purified VLPs were mixed with excipients sorbitol (10mM or 1.2% w/v) polysorbate 80 (PS80; 0.076mM or 0.01% w/v), L-histidine (10mM or 0.16% w/v) in concentrations used in the commercial vaccines MMR and HPV. The stability of the VLP mixtures incubated at 4°C for up to 14-days was analysed using several methods. First, we determined the antigenicity of purified ß-SEM-SARS-CoV-2 VLPs by ELISA probing with a polyclonal anti-S antibody. The antigenicity of VLPs in PBS alone waned after 7 days and by day 14 were weakly reactive (Figure 2A). VLPs mixed with PS80 appeared to be more stable with only a small decrease in antigenicity between days 7 and 14. In contrast, the antigenicity of VLPs mixed with sorbitol and L-histidine remained stable to 14 days, with VLPs in sorbitol exhibiting the best stability (Figure 2 C and D).

**Figure 2.**
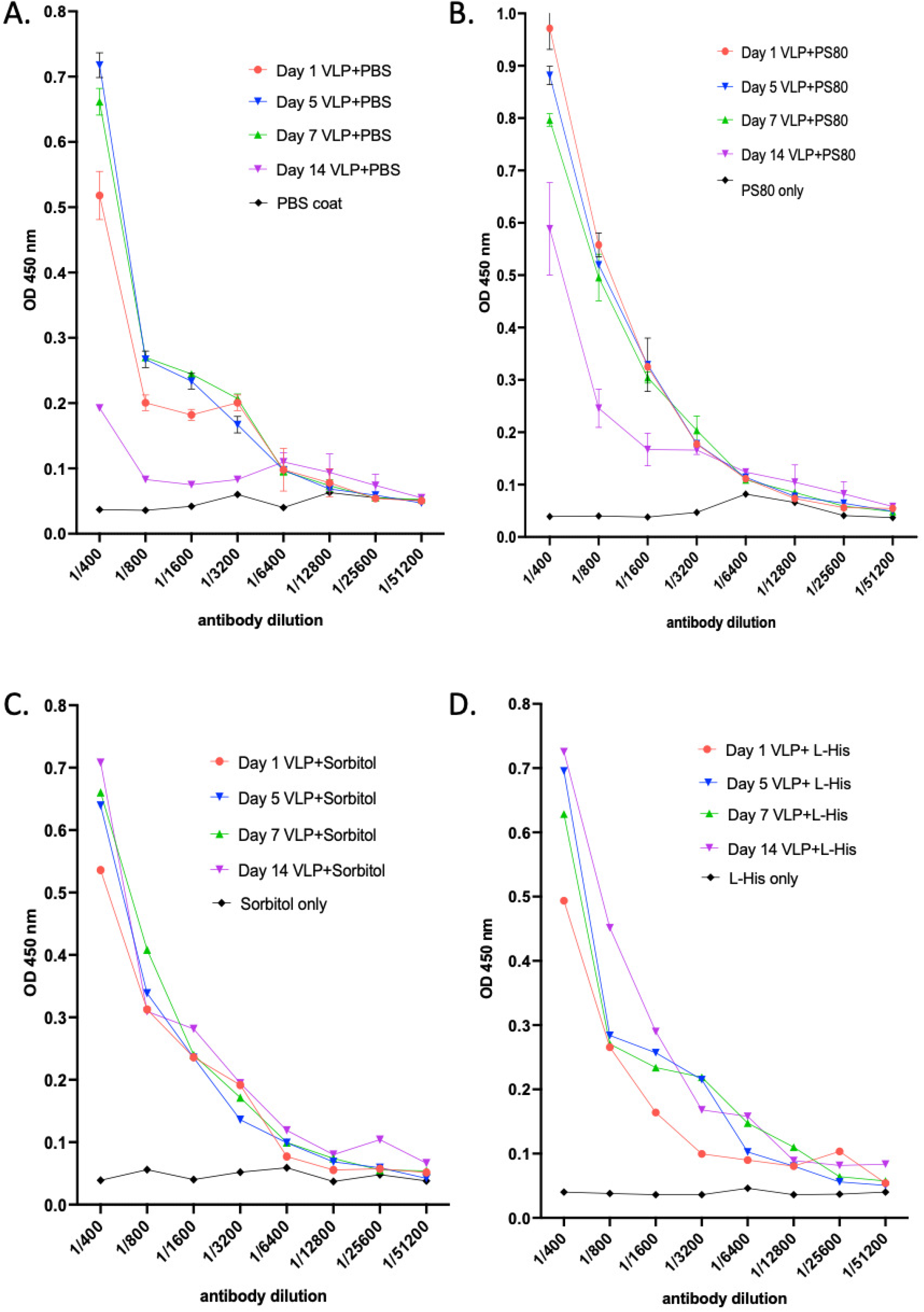
ELISA analysis of ß-SARS-CoV-2 VLPs in PBS or mixed with excipients PS80, sorbitol or L-histidine and incubated at 4 °C for 14 days. Plates were coated with ß-SARS-CoV-2 VLPs followed by probing with anti-Spike antibody in 2-fold dilutions from 1:400 to 1:51200 (mean ± SD). ELISAs were performed three times and each dilution tested in triplicate. Samples were collected on days 1, 5, 7 and 14 for ELISA analysis, probing with polyclonal anti-S antibody. Experiments were performed in triplicate.

### 3.2 Western immunoblot of ß-SARS-CoV-2 VLPs

The composition of the ß-SARS-CoV-2 VLPs mixed with sorbitol, polysorbate 80 or L-histidine and incubated for 14-days at 4°C was determined by Western immunoblot probing with anti-S, anti-E and anti-M antibodies. Spike, membrane and envelope proteins were detected in all samples from VLPs mixed with PBS, PS80, sorbitol and L-histidine and remained unaltered in size from day 1 through day 14 (Figure 3). The spike protein retained a consistent size of 115kDa thorough the 14-day period. Similarly, there was no change in the size of the membrane protein (25kDa) from day 1 to day 14. The E protein was detected predominantly in its dimeric form (24kDa), consistent with previous reports (21, 23) and like the spike and membrane proteins did not change in size over the 14 days.

**Figure 3.**
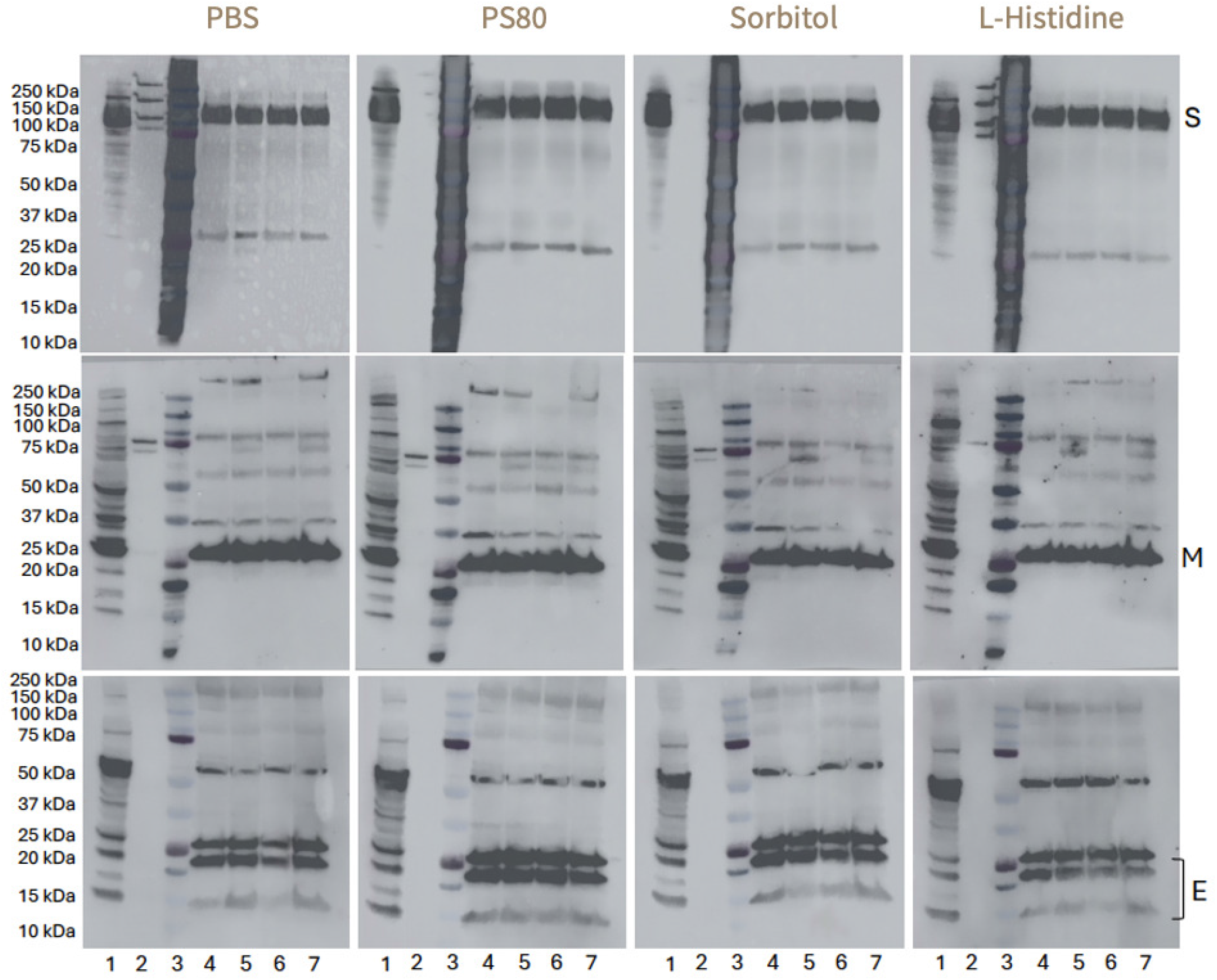
Western immunoblot of purified ß-SARS-CoV-2 VLPs in PBS or mixed with PS80, sorbitol or L-histidine and incubated at 4°C for 14 days. Immunoblots were probed with polyclonal antibodies against the viral spike, membrane and envelope proteins. Lane 1: Ancestral-SEM VLP crude extract; Lane 2: Uninfected Vero cell supernatant; Lane 3: Biorad Dual Colour standard; Lane 4: βS_13_EM_sh_ Batch 13 Day 1; Lane 5: βS_13_EM_sh_ Batch 13 Day 5; Lane 6: βS_13_EM_sh_ Batch 13 Day 7; Lane 7: βS_13_EM_sh_ Batch 13 Day 14.

### 3.3 Dynamic light scatter analysis of ß-SARS-CoV-2 VLPs

Having shown that the spike, membrane and envelope proteins were maintained through the 14 days of incubation, we next determined whether the VLPs retained their particulate structure. We utilized dynamic light scatter (DLS) to evaluate the size and distribution of purified ß-SARS-CoV-2 VLPs in PBS or mixed with excipients PS80, sorbitol or L-histidine (11). The DLS analysis showed a polydisperse distribution characterized by two distinct peaks at 183 ± 6.8 nm and 12 ± 3 nm with the larger peak consistent with intact VLPs and the smaller peak with the presence of free spike trimers as we have previously reported (11) (Figure 4A). The VLPs maintained their dimensions and particulate structure through the 14-day incubation, and this finding was consistent with VLPs in PBS or mixed with stabilising excipients, with little difference observed between VLPs in PS80, sorbitol or L-histidine (Figure 4). In addition, the polydispersity index for all samples was below 0.1, indicating a high degree of uniformity of VLPs regardless of the excipient used. Finally, purified ß-SARS-CoV-2 VLPs in PBS and stored for 24 months at -80°C retained their DLS profile and size distribution (Figure 4 B).

**Figure 4.**
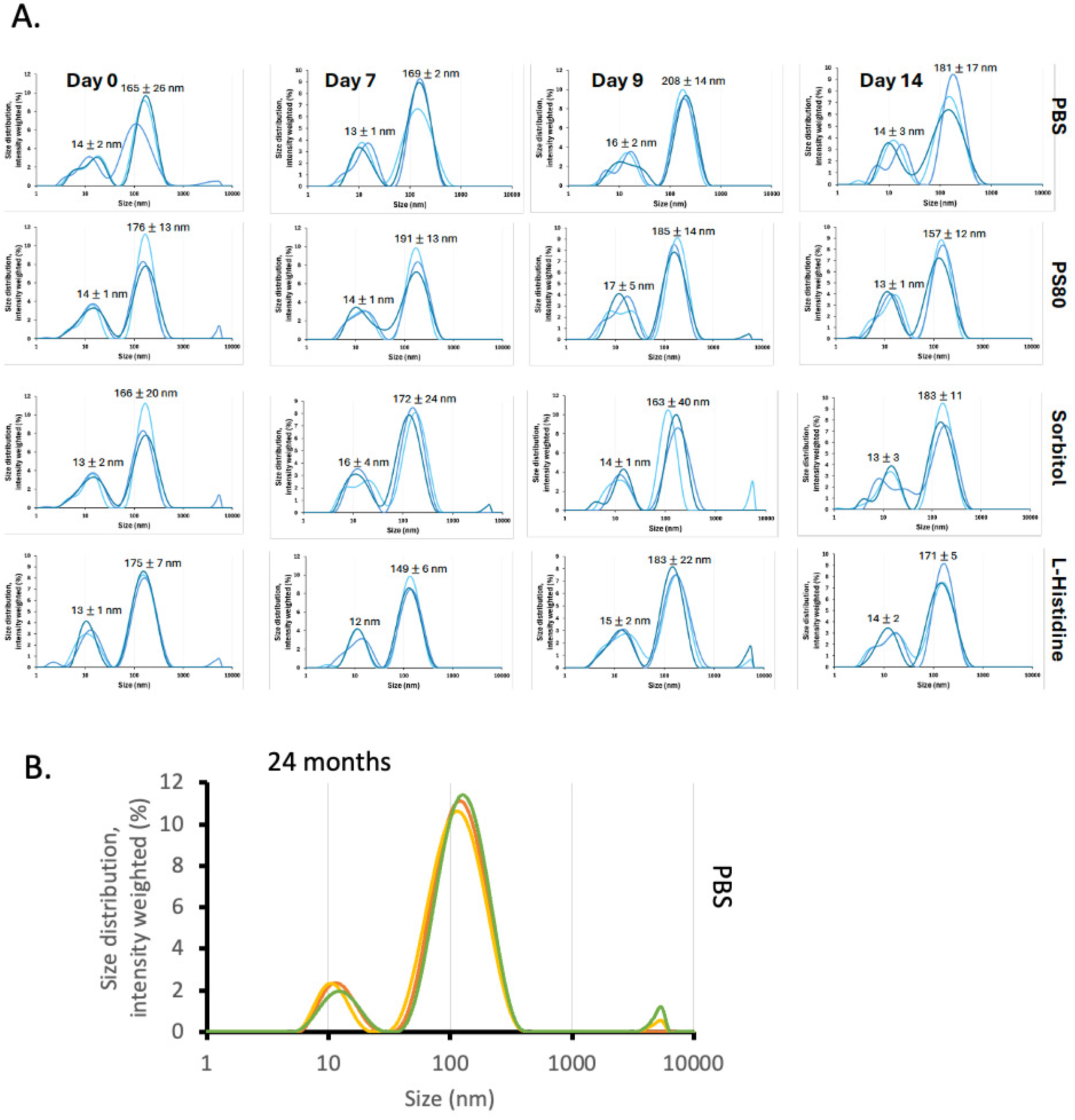
Dynamic light scatter analysis. (A) Purified ß-SARS-CoV-2 VLPs in PBS or mixed with PS80, sorbitol or L-histidine were incubated at 4 °C for 14 days and samples collected at days 7, 9 and 14 for DLS analysis. Two distinct peaks were identified in all excipient groups, the size of peaks are consistent with Spike trimers (left, smaller peak) and ß-SARS-CoV-2 VLPs (right, larger peak). (B) Purified ß-SARS-CoV-2 VLPs in PBS and stored for 24 months at -80°C. Experiments were performed in triplicate.

## 4. Discussion

In this study we have shown that our SARS-CoV-2 VLP vaccine is stable at 4°C for up to 4 months and the addition of stabilizing excipients improves stability when compared to VLPs incubated in PBS alone. The addition of excipients helped to maintain the immunogenicity of the VLPs as shown by ELISA. DLS analysis revealed that the VLPs retained their particulate structure regardless of whether they were stored in PBS or with excipients. In addition, the particles retained their size distribution and remained stable for up to 2 years when stored at - 80°C. Our results are encouraging for showing that our SARS-CoV-2 VLPs are stable for extended periods but further studies using combinations of excipients and more comprehensive analysis of thermal stability will be needed to properly characterise vaccine quality and its shelf-life through the various stages of the product life cycle. These data are essential to progress the vaccine to clinical trials, for licensure and to provide a methodological process that can be applied to test future batches of manufactured vaccine (24).

Different types of stabilizers also have important roles in maintaining the integrity of a VLP based vaccine. Non-ionic surfactants like polysorbate 80 are important to help prevent protein based vaccines adsorbing to surfaces like vials as well as inhibiting the proteins from aggregating, denaturing or precipitating (16). Non-ionic surfactants like Polysorbate 80 (PS80), Triton-X 100 and 114, Brij 35 and 58 at concentrations of 0.01% have all been shown to enhance stability and prevent aggregation of HPV VLPs. Even concentrations as low as 0.001% of PS80 enhanced storage stability of HPV VLPs (16). To achieve optimal stabilization of the HPV VLPs a minimum or physiological salt concentration (e.g. 150mM NaCl) is required (16), although the addition of PS80 allows for lower salt concentrations to be used (25). We used PS80 at a final concentration of 0.01% with a physiological salt concentration of 137mM and found that this improved stability of VLPs and prevented aggregation as shown by DLS analysis. Higher concentrations of PS80 and NaCl have also been shown to protect HPV VLP integrity against the effects of freeze-thawing and will warrant further investigation with our ß-SARS-CoV-2 VLPs (25).

Similarly, sugars like sorbitol, sucrose, mannitol and trehalose and amino acids like L-histidine, proline, glycine, glutamate and aspartate can stabilize VLPs from the effects of freeze-thawing and freeze drying (15) (14, 16). The commercial measles, mumps, rubella (MMR) vaccine contains sorbitol in a final concentration of 2.9% (w/v) while the HPV vaccine contains 0.156% (w/v) L-histidine. We used a sorbitol concentration of 1.2% (w/v) and 0.156% (w/v) L-histidine which, like PS80, maintained the stability and integrity of the VLPs. Analysis by DLS also showed that the VLPs retained their size and did not aggregate.

The stabilizing effects of amino acids has also been more clearly elucidated in a study that showed that amino acids adsorb onto the surface of proteins, enhancing protein stability. The amino acid-protein interaction is also more effective for proteins of opposite charge (26). For a complex structure like a VLP for which determining overall charge may not be possible, stabilization may be better achieved by using combinations of amino acids of different charge. The inclusion of combinations of amino acids to improve our VLP vaccine stability will require more comprehensive analyses.

We used DLS to provide a measure of particle size and aggregation and showed that the VLPs retained their size and integrity throughout a 14-day incubation at 4°C but also for up to 2 years stored at -80°C. This finding is encouraging, however, more comprehensive information about the characteristics of our VLP vaccines could be provided with techniques such as asymmetrical flow field flow fractionation and electrospray differential mobility analysis (15, 20, 27) that may provide more information on the VLP characteristics. These analyses would provide more comprehensive data to support regulatory submissions and of the ability of VLPs stored under these conditions to stimulate an appropriate immune response and will warrant further investigation with our vaccine.

## Author contribution statement

**Melissa Edeling:** Writing – _review & editing, Writing – _original draft, Visualization, Methodology, Formal analysis, Data curation. **Linda Earnest:** Writing – _review & editing, Methodology, Formal analysis, Data curation. **Julio Carrera Montoya:** Writing – _review & editing, Validation, Methodology, Formal analysis, Data curation. **Ashley Huey Yiing Yap:** Writing – _review & editing, Methodology, Formal analysis, Data curation. **Dhiraj Hans:** Writing – _review & editing, Project administration, Conceptualization. **Joseph Torresi:** Writing – _original draft, Writing – _review & editing, Visualization, Validation, Supervision, Project administration, Methodology, Formal analysis, Conceptualization.

## Acknowledgements

This work was supported by grants from the National Health and Medical Research Council (NHMRC) Medical Research Future Fund (MRFF)

## Data availability

Data will be made available on request.

